# Proteasome inhibition induces DNA methylation alteration by attenuating the synthesis of DNA methyltransferase 1 and 3B in colorectal cancer

**DOI:** 10.1101/2024.06.14.598995

**Authors:** Wenwen Zhou, Yuling Sheng, Dingxue Hu, Yunyun An, Mengqi Yang, Wanqiu Wang, Shiva Basnet, Jingyu Yan, Shuxia Zhang, Qi Liu, Yunze Li, Yi Tan, Jing Gao, Kun Sun, Changzheng Du

## Abstract

Proteasome is an essential organelle in guarding cellular protein homeostasis. Here, we report that inhibition of proteasome leads to alterations in DNA methylation patterns in colorectal cancer (CRC) by surpressing the synthesis of DNA methyltransferases (DNMTs). We found that treating CRC cells with proteasome inhibitors results in attenuated translation of DNMT1 and DNMT3B, mediated by the inactivation of AKT and mammalian target of rapamycin (mTOR), which is dependent on the accumulation of p300, an acetyltransferase that inhibits AKT through acetylation modification. Furthermore, we demonstrated that downregulation of DNMT1 and DNMT3B confers protection against proteasome inhibitor treatment, potentially through reprogramming the transcriptome of CRC cells, highlighting the significant role of DNMTs in response to disruptions in protein homeostasis. Interestingly, the proteasome inhibitor-induced downregulation of DNMT1 and DNMT3B appears to be CRC specific, notwithstanding the underlying mechanism remains unclear. Altogether, our findings reveal an epigenetic effect of proteasome on DNA methylation in CRC through its regulation of DNA methyltransferase synthesis.

## Introduction

Proteasome is the major organelle responsible for ubiquitinated protein degradation in eukaryotic cells, with cellular functions ranging from general protein homeostasis to stress response and cell survival (Bard et al, 2018; Rousseau & Bertolotti, 2018). Inhibition of proteasome leads to accumulation of misfolded or deleterious proteins, triggering complicated biological effects and toxicity to cells (Thibaudeau & Smith, 2019); therefore, numerous proteasome inhibitors (PIs) have been developed as anti-cancer reagents for clinical treatment (Fricker, 2020). However, the pharmacological mechanism of PIs in cancer treatment is not fully understood; moreover, due to the complicated pharmacological effects of PIs, it remains unclear which phenotype of cancer is sensitive to PI therapy in clinical practice.

In addition to disrupting proteostasis, inhibition of proteasome also leads to transcriptional and epigenetic changes, including alterations in histone modification and gene expression, by targeting epigenetic regulators (Kamens et al, 2023; Kinyamu et al, 2020). In the present study, we focused on the epigenetic impact of PIs on DNA methylation due to its critical role in carcinogenesis and clinical treatment in cancer (Nishiyama & Nakanishi, 2021). In mammals, DNA methylation is catalyzed by DNA methyltransferases (DNMTs), which are frequently mutated or overexpressed in cancer (Greenberg & Bourc’his, 2019; Zhang et al, 2020b), and is counteracted by demethylases such as Tet methylcytosine dioxygenases (TETs) (Greenberg & Bourc’his, 2019). DNMT family is a target for cancer therapy, with numerous DNMT inhibitors currently undergoing clinical trials (Mehdipour et al, 2021; Nishiyama & Nakanishi, 2021). Our investigation into DNA methylation in colorectal cancer (CRC) revealed distinct changes following PI treatment. We further revealed that PIs induced a remarkable downregulation of DNMT1 and DNMT3B in CRC, contrary to the reported accumulation observed in other types of cancer (Leng et al, 2018). Moreover, we demonstrated that the PI-induced downregulation of DNMT1/3B is mediated by p300 accumulation and subsequent AKT-mTOR pathway inactivation, leading to a translational inhibition of DNMT1/3B. Functionally, our findings suggest that downregulation of DNMT1/3B protects CRC cells from PI treatment, potentially by reprogramming the transcriptome of CRC.

## Results and discussion

### Proteasome inhibition alters DNA methylation profile in CRC

To assess the impact of proteasome inhibition on DNA methylation in CRC, we utilized the Infinium Methylation EPIC v2.0 BeadChip which covers over 900,000 CpG probes across the human genome (Noguera-Castells et al, 2023). CRC cells were continuously treated with MG132 (0.2μM) for 21 passages, and DNA methylation levels were assessed at passages 14 and 21, with DMSO-treated cells used as a control. Our results revealed a significant increase of the CpG sites with altered methylation across the genome as the treatment prolonged (Fig. 1A-B); and interestingly, there was a large number of demethylated CpG sites showed up at passage 21 compared to passage 14 (Fig. 1B). These findings led us to further investigate the alteration of the enzymes responsible for maintaining DNA methylation in CRC.

**Figure 1.**
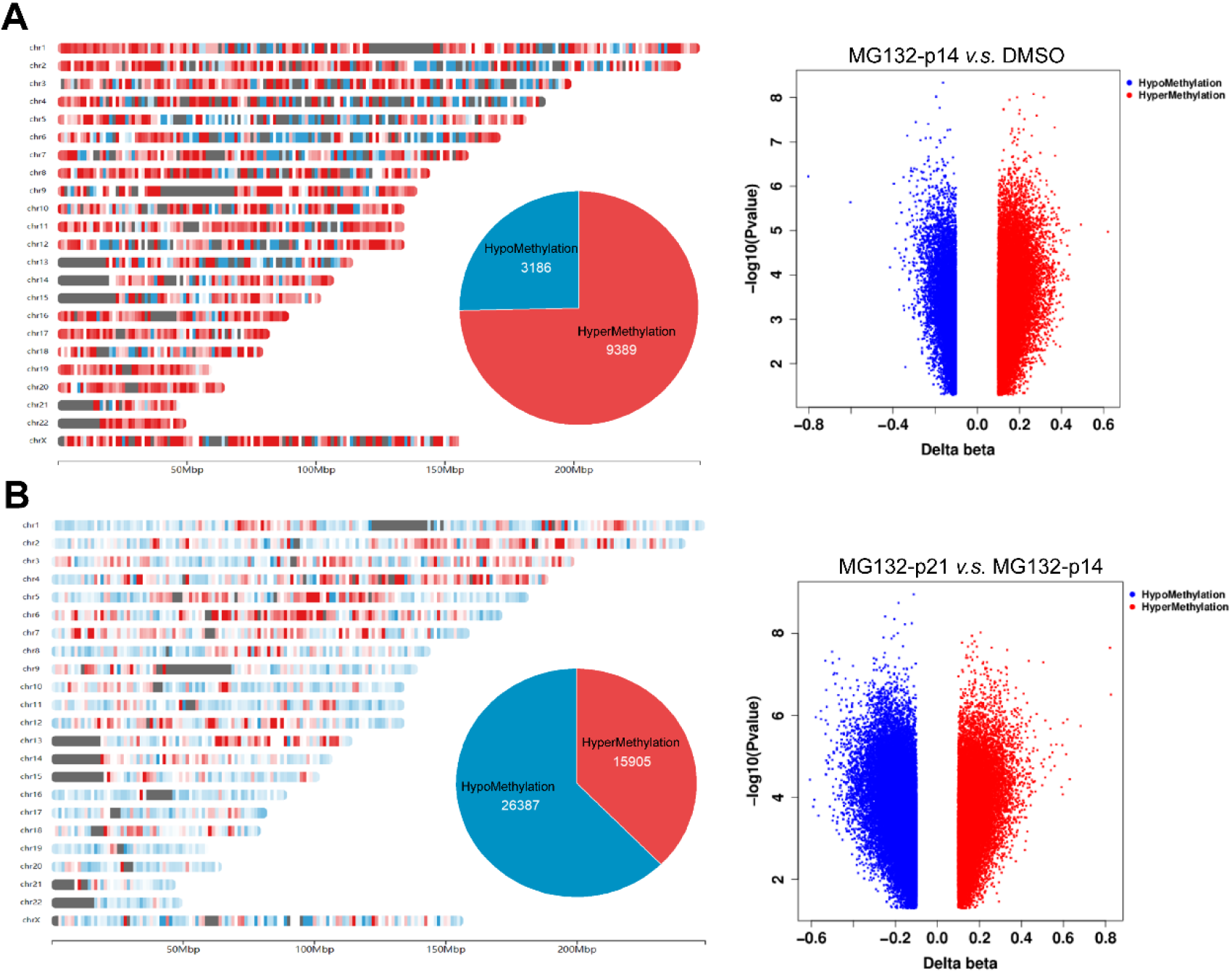
Proteasome inhibition alters DNA methylation profile in colorectal cancer. **(A).** Chromosomal distribution of hypo- or hyper-methylation regions in the passage 14 of HT29 cells continuously treated by MG132 (left panel), and the volcano plots of altered methylation probes (right panel), using DMSO treated cells as control. **(B).** Chromosomal distribution of hypo- or hyper-methylation regions in the passage 21 of HT29 cells continuously treated by MG132 (left panel), and the volcano plots of altered methylation probes (right panel), using passage 14 cells as control.

### Proteasome inhibition attenuates translation of DNMT1 and DNMT3B by inactivating mTOR in colorectal cancer cells

Given that DNA methylation is catalyzed and maintained by DNMTs, and waived by demethylation enzymes such as TET family in mammals (Greenberg & Bourc’his, 2019; Zhang et al, 2020b), we initially detected the abundance of DNMTs and TETs. We found that MG132 induced a remarkable downregulation of DNMT1 and DNMT3B, which was dose- and time-dependent (Fig. 2A-B), without affecting DNMT3A and TET family (Fig. S1A). The downregulation of DNMT1 and DNMT3B was also induced by other PIs (Fig. 2C-D); however, this phenotype was not observed in a handful of non-CRC cell lines examined (Fig. S1B). Given that PIs could activate autophagy (Pohl & Dikic, 2019; Wang et al, 2019), we blocked autophagy using chloroquine, a commonly used lysosomal inhibitor, finding that blocking autophagy did not rescue the MG132-induced downregulation of DNMT1 and DNMT3B (Fig. 2E), ruling out the possibility of antophagy-dependent protein degradation. Additionally, there was no change in the transcription of DNMT1 and DNTM3B following PI treatment as determined by RT-qPCR assay (Fig. S1C-D).

**Figure 2.**
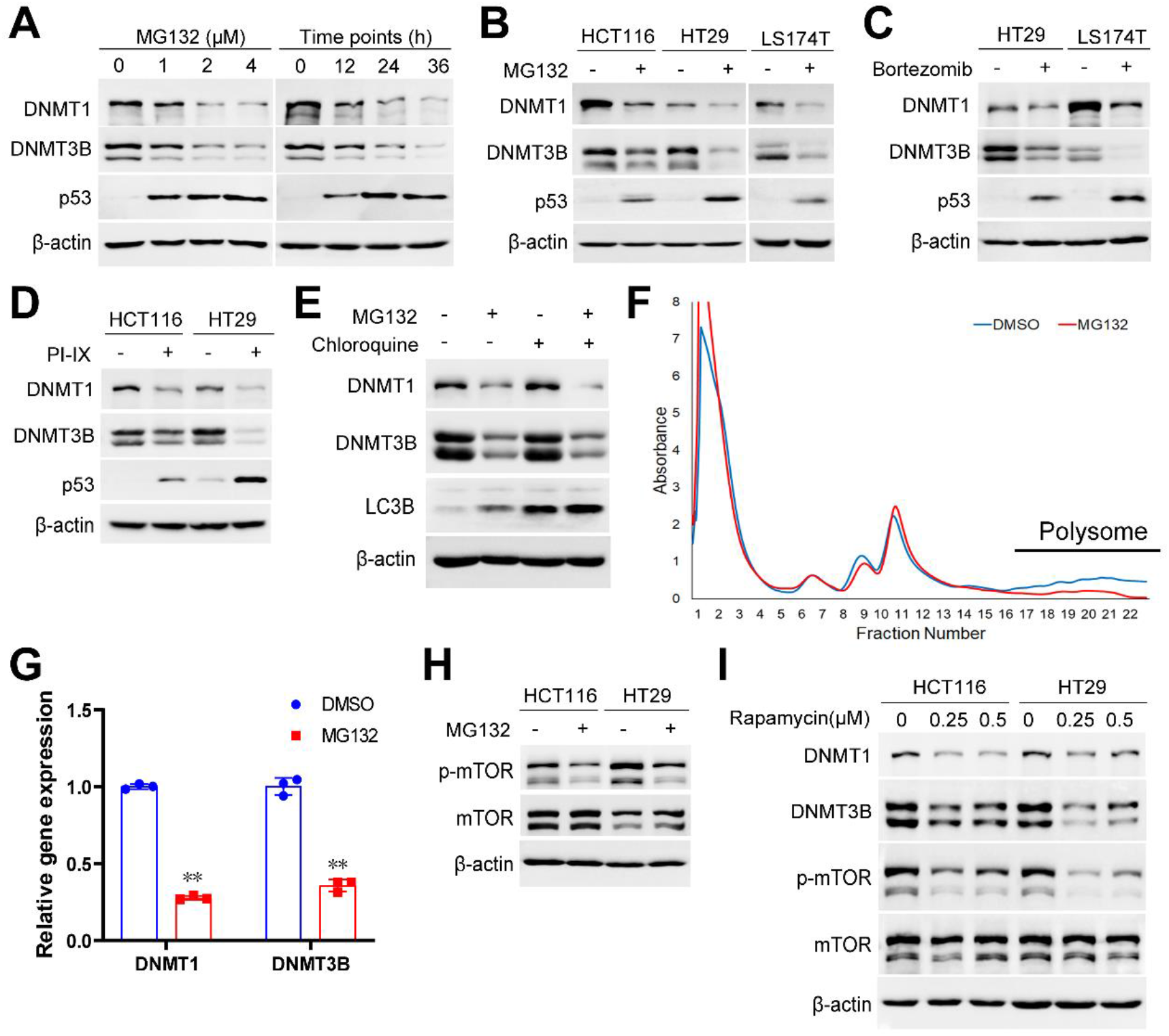
Proteasome inhibition blocks the translation of DNMT1 and DNMT3B. **(A).** The abundance of DNMT1 and DNMT3B following the treatment of MG132 with different doses (left) and time-points (right) in HT29 cells were detected by immunoblotting, using p53 as a positive control. **(B).** The abundance of DNMT1 and DNMT3B following the treatment of MG132 (1μM for 24 hours) in HCT116, HT29 and LS174T cells were detected by immunoblotting. **(C-D)**. The abundance of DNMT1 and DNMT3B following the treatment of Bortezomib (100nM for 24 hours) (C) or PI-IX (10μM for 24 hours) (D) in multiple CRC cells were detected by immunoblotting. **(E).** HT29 cells without or with MG132 treatment were treated with DMSO or Chloroquine of 20μM for 24 hours, the abundance of DNMT1 and DNMT3B was determined by immunoblotting. **(F).** Polysome profiling analysis was performed in HT29 cells treated with DMSO or MG132 (1μM for 24 hours), and polysomal RNA (as labeled) was collected and reversely transcripted. **(G).** The mRNA abundance of DNMT1 and DNMT3B in the polysomal RNA was detected by RT-qPCR assay. **(H).** The phosphorylation of mTOR (Ser2448) in HT29 without or with MG132 treatment (1μM for 24 hours) were detected by immunoblotting. **(I).** The abundance of DNMT1 and DNMT3B without or with treatment of Rapamycin (24 hours) were detected by immunoblotting. \*\**P*<0.01

To investigate whether PIs inhibit the translation of DNMT1/3B, we performed polysome profiling assay, finding a global inhibition of protein translation induced by PIs (Fig. 2F); furthermore, the DNMT1/3B specific translation also displayed a significant decrease, identified by RT-qPCR assay using the polysomal RNA (Fig. 2G).

As a key regulator of metabolism and protein translation, mTOR is significantly activated in cancer (Glaviano et al, 2023; Panwar et al, 2023). When protein degradation is blocked by PIs, mTOR rapidly responds by reducing the phosphorylation of downstream translation factors including p70 S6 kinase and 4E-BP1 (Battaglioni et al, 2022; Cao et al, 2023), meanwhile activating autophagy through the dephosphorylation of its downstream autophagy proteins ULK1/ATG13/FIP200 complex (Dossou & Basu, 2019), to maintain intracellular proteostasis. Based on the reports that PIs repress the activity of mTOR (Driessen et al, 2018; Paradzik et al, 2021), we detected the phosphorylation of mTOR, validating that MG132 treatment reduced the phosphorylation of mTOR (Fig. 2H). To verify the role of mTOR in the translational regulation of DNMT1/3B, we treated CRC cells with rapamycin, a mTORC1-specific inhibitor (Panwar et al, 2023), revealing that rapamycin downregulates the DNMT1 and DNMT3B, in agreement of what was observed following PIs treatment (Fig. 2I). Above all, these data support the notion that PIs inhibit protein translation of DNMT1 and DNMT3B by inactivating mTORC1 in CRC.

### Proteasome inhibition inactivates AKT-mTOR signal pathway by accumulation of p300

Given the crucial role of AKT as an upstream kinase in activating mTOR (Glaviano et al, 2023; Panwar et al, 2023), we detected the alteration of AKT phosphorylation following PI treatment in CRC cells, finding that PIs inhibit AKT by inducing its dephosphorylation at S473 (Fig. 3A). Subsequently, we discovered that inhibition of AKT using its inhibitor BAY1125976 led to a significant decrease of mTOR phosphorylation and downregulation of DNMT1/3B (Fig. 3B), reminiscent of what was observed in MG132 treated CRC cells. These results collectively support the notion that the PI-induced translational blockage of DNMT1/3B is mediated by AKT inactivation. Among the upstream regulators affecting AKT phosphorylation, the histone acetyltransferase p300 (KAT3B) deems to be a potential mediator of PI-induced AKT inactivation for two reasons: 1. p300 inhibits the phosphorylation of AKT by mediating its acetylation at K20 (Sundaresan et al, 2011), while this acetylation in the PH domain of AKT blocked the binding of AKT to PIP_3_, preventing its membrane localization and phosphorylation (Sundaresan et al, 2011). 2. The degradation of p300 is ubiquitin-dependent and proteasome-mediated in cancer cells (Xu et al, 2020), suggesting the PI treatment potientially increase its abundance in CRC. Therefore, we hypothesize that PIs inactivate AKT and mTOR through accumulation of p300 in CRC cells. A number of complementary findings supported our hypothesis: First, we confirmed the accumulation of p300 protein following MG132 treatment in CRC cells (Fig. 3C). Second, we demonstrated that overexpressing exogenous p300 led to significant dephosphorylation of AKT at Ser 473, resulting in downregulation of DNMT1/3B (Fig. 3D); complementarily, knockdown of p300 promotes AKT phosphorylation and DNMT1/3B upregulation (Fig. 3E). Moreover, knockdown of p300 reversed the

**Figure 3.**
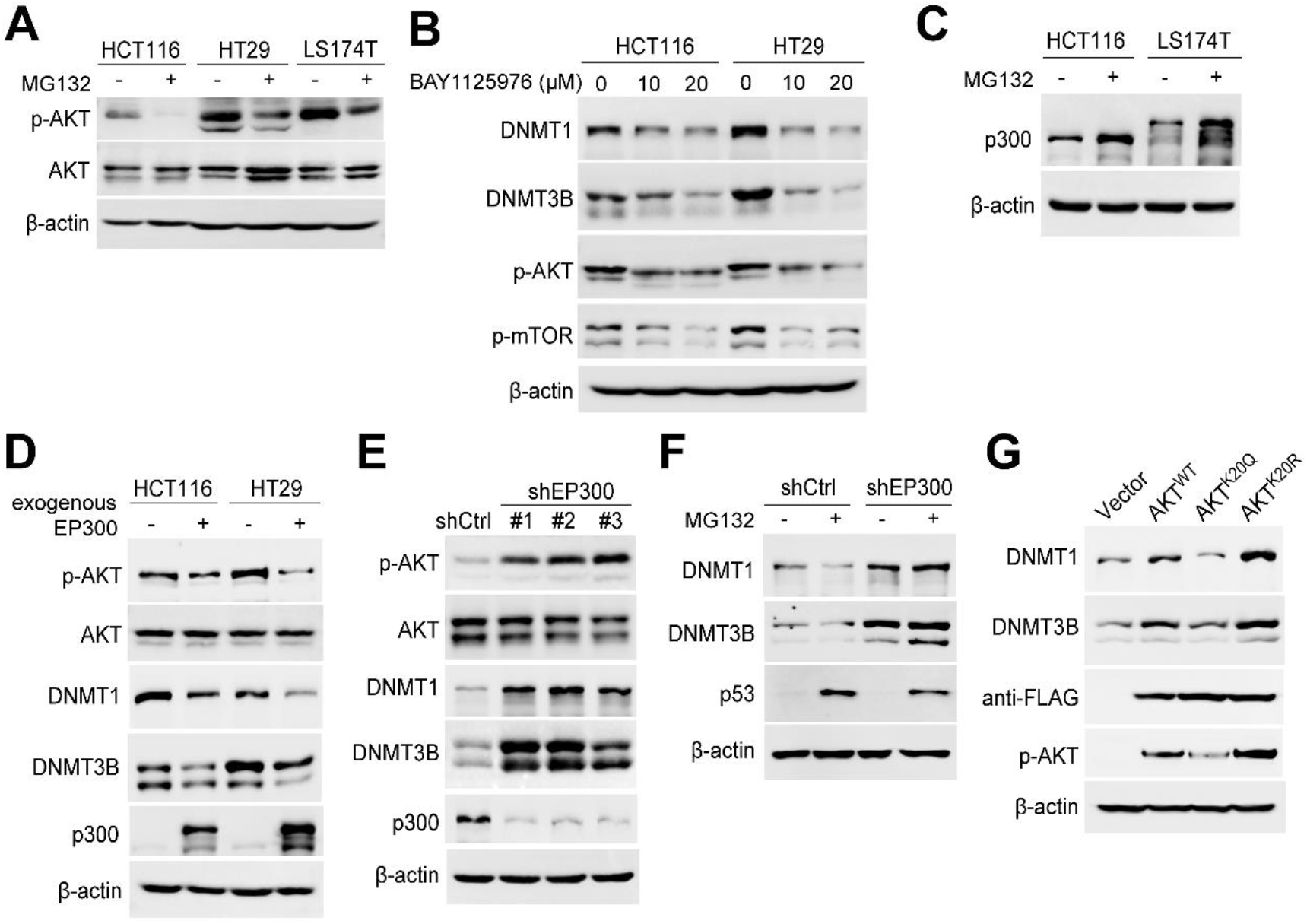
Proteasome inhibition inactivates AKT-mTOR signal pathway by accumulation of p300. **(A).** The phosphorylation of AKT (Ser473) in HCT116, HT29 and LS174T without or with treatment of MG132 (1μM for 24 hours) was detected by immunoblotting. **(B).** The abundance of DNMT1 and DNMT3B in HCT116 and HT29 without or with treatment of BAY1125976 (10 or 20μM for 24 hours) was detected by immunoblotting, using p-AKT and p-mTOR as positive controls. **(C).** The abundance of p300 in HT29 without or with treatment of MG132 (1μM for 24 hours) was detected by immunoblotting **(D).** The phosphorylation of AKT (Ser473) and the abundance of DNMT1 and DNMT3B without or with exogenous p300 overexpression in HCT116 and HT29 cells was determined by immunoblotting. **(E).** The phosphorylation of AKT (Ser473) and the abundance of DNMT1 and DNMT3B without (shCtrl) or with p300 knockdwon (shEP300) in HT29 cells was determined by immunoblotting. **(F).** HT29 derivative cell lines (shCtrl or shEP300) were treated with DMSO or MG132 (1μM for 24 hours), the abundance of DNMT1 and DNMT3B was determined by immunoblotting. **(G).** The abundance of DNMT1 and DNMT3B in the HT29 cells, without (Vector) or with exogenous expression of wild-type, K20Q or K20R mutant AKT (FLAG-tagged), was determined by immunoblotting.

MG132-induced downregulation of DNMT1/3B (Fig. 3F), indicating that p300 mediates the PI-induced DNMT1/3B downregulation. Finally, we generated mutant AKT variants on its acetylated lysine K20 (K20Q and K20R), to validate the role of acetylation in AKT phosphorylation and DNMT1/3B abundance. We found that the acetylation-mimic mutation (K20Q) attenuated its phosphorylation at S473, and downregulated DNMT1 and DNMT3B; in contrast, the acetylation-dead mutation (K20R) promoted its phosphorylation and upregulated DNMT1 and DNMT3B (Fig. 3G). Taken together, these data supported the notion that PIs inactivate AKT-mTOR signaling pathway through accumulation of p300.

To illuminate the mechanism of the CRC-specific downregulation of DNMT1/3B, we detected p300 and AKT-mTOR activity in non-CRC cells. Interestingly, we observed that PIs could induce the accumulation of p300 and the inactivation of AKT and mTOR in the MCF-7 breast cancer cell line (Fig. S2A). However, inhibition of mTORC1 using rapamycin did not induce a downregulation of DNMT1/3B in MCF-7 (Fig. S2B), suggesting that mTORC1 does not regulate the translation of DNMT1/3B in this cell line. Furthermore, we investigated other potential regulators involved in PI-induced AKT-mTOR pathway inactivation, such as DEPTOR which is known to be upregulated after MG132 treatment in myeloma (Vega et al, 2022), our immunoblot data revealed that DEPTOR was not accumulated following MG132 treatment in CRC cells (Fig. S2C). Additionally, considering previous reports suggesting that PIs inhibit AKT by enhancing PP2A activity (Wei & Xia, 2006); we treated CRC cells with PP2A inhibitor Okadaic acid (OA) and found no alteration in DNMT1/3B abundance (Fig. S2D), thereby excluding PP2A as a mediator for the PI-induced downregulation of DNMT1/3B.

### Downregulation of DNMT1 and DNMT3B protects cancer cells against proteasome inhibition in CRC

To identify the role of PI-induced DNMT1/3B downregulation, we conducted a double-knockdown of DNMT1 and DNMT3B (shDNMT1&3B) in CRC cells to mimic the PI-induced downregulation (Fig. 4A). Our results demonstrated that simultaneous knockdown of DNMT1 and DNMT3B significantly enhances the cellular resistance to PI treatment, as determined by CCK8 assay (Fig. 4B-C) and clone formation assay (Fig. 4D-E). To reveal the underlying mechanism, we individually knocked down DNMT1 (shDNMT1) and DNMT3B (shDNMT3B) respectively, and found a significant alteration of transcriptome following depleting either DNMT1 (shDNMT1) or DNMT3B (shDNMT3B), with more upregulated genes than downregulated genes in each knockdown cells (Fig. 4F). Interestingly, knockdown of either DNMT resulted in the downregulation of specific core components of the proteasome machinery (Fig. 4G-H), which are also the targets of PIs (Larsson et al, 2022; Yadav et al, 2023), potentially reducing vulnerability to PIs. Above all, these data support the protective role of DNMT1 and DNMT3B downregulation against proteasome inhibition in CRC through modulation of the cellular transcriptome.

**Figure 4.**
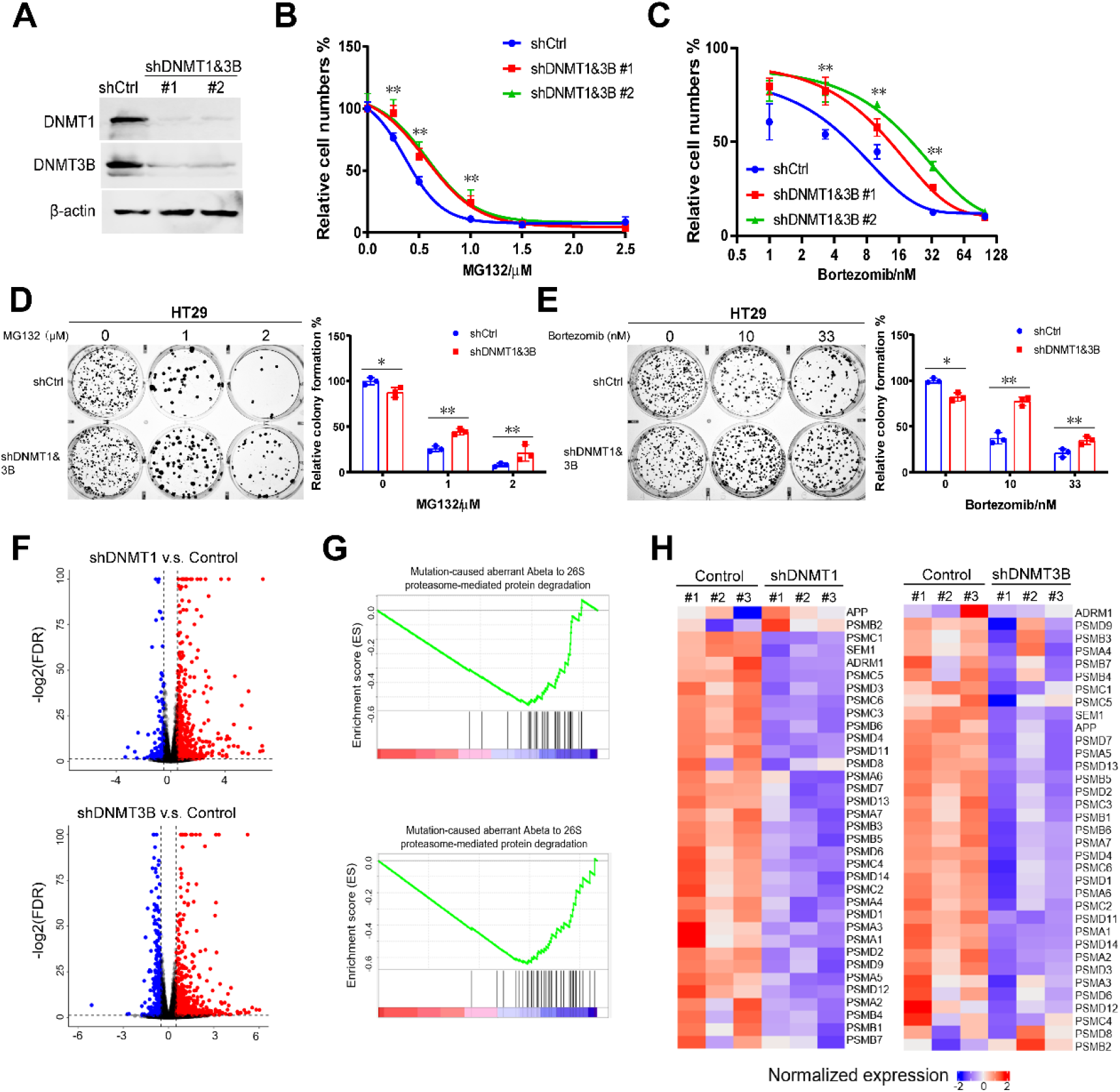
Downregulation of DNMT1 and DNMT3B protects cancer cells against proteasome inhibition. **(A).** The abundance of DNMT1 and DNMT3B in the HT29 cells without (shCtrl) or with double-knockdown of DNMT1 and DNMT3B (shDNMT1&3B), detected by immunoblotting. **(B-C).** HT29 derivative cell lines (shCtrl or shDNMT1&3B) were treated with MG132 (B) or Bortezomib (C) for 48 hours, respectively; and cell numbers were determined by CCK8 assay. **(D-E).** HT29 derivative cell lines (shCtrl or shDNMT1&3B) were treated with MG132 (D) or Bortezomib (E) for 24 hours, and colony formation assays were performed with 500/well to determine the proliferation ability of treated cells. **(F).** Volcano plots of different expressed genes (DEGs) in HT29 cells without (shCtrl) or with knockdown of DNMT1 (upper) or DNMT3B (bottom). **(G).** Functional enrichment analysis of DEGs in HT29 cells without (shCtrl) or with knockdown of DNMT1 (upper) or DNMT3B (bottom), within which, the proteasomal degradation pathway was enriched. **(H).** Heatmap of DEGs enriched in proteasomal degradation pathway.

In conclusion, this study identified an epigenetic effect of proteasome inhibition that it induces a translational inhibition of DNA methyltransferases and an alteration of DNA methylation profile in colorectal cancer.

## Methods

### Cell culture

All cell lines in this study were purchased from American Type Culture Collection (ATCC, USA). The human CRC cell lines HT-29, LS174T and Colo320 were cultured in Roswell Park Memorial Institute (RPMI) 1640 medium (Bio Basic Cat# E600028), and HCT116, SW48, SW620 were cultured with Dulbecco’s Modified Eagle Medium (DMEM) (Gibco Cat# C11965500BT; the human embryonic kidney cell line HEK-293T, human breast cancer cell line MCF-7, glioblastoma cell line U-251MG, gastric cancer cell line SNU-1 and pancreatic cancer cell line PANC-1 were cultured in DMEM. All the culture mediums were supplemented with 10% fetal bovine serum (FBS, Procell Cat# 164210), penicillin (100 U/mL) and streptomycin (100μg/mL). All cell lines were cultured at 37°C in a humidified atmosphere of 5% CO_2_.

### DNA extraction, bisulfite conversion and DNA methylation microarray

CRC Cells were harvested and lysed with lysis buffer (10 mM Tris pH 8.0, 300 mM NaCl, 5 mM EDTA pH 8.0 and 0.5% SDS), and genomic DNA was extracted using as previously reported (Jiang et al, 2020). Bisulfite conversion of DNA was performed using the EZ DNA methylation kit (Zymo Research, Cat# D5001) following its manufactural protocol. DNA methylation were detected and evaluated using Infinium MethylationEPIC v2.0 Kit as previously reported (Noguera-Castells et al, 2023). The DNA methylation assay was repeated 3 times for each condition: DSMO control, p14 and p21 after PI-treatment. Methylation densities of each CpG site were compared using t-tests between two conditions (i.e., p14 vs DSMO, and p21 vs p14) and the p-values were corrected using Benjamini and Hochberg method. CpG sites with adjusted p-values lower than 0.05 and differences in methylation densities larger than 10% were considered as differentially methylated.

### Cell viability and proliferation assays

Cell viability following the PI treatment was assessed using Cell Counting kit-8 (CCK-8, Beyotime, Cat# C0040) according to the manufacturer’s instruction. Briefly, cells were seeded in 96-well plates at 2×10^4^ cells per well. After culturing for 12 hours, cells were treated with different concentration gradients of PIs for 48 hours; then CCK8 solution was added into medium at a concentration of 10% (v/v), incubating in 37°C for 1 hours before measurement. Absorbance of each well was determined at 450 nm using a Synergy HTX microplate reader (Agilent, USA). The relative cell numbers were calculated and presented as mean value ± standard deviation (mean ± SD).

Cell proliferation was assessed by colony formation assay. Briefly, cells plated at a confluence of 70%-80% were treated with PIs for 24 hours, then seeded on 6-well plates at a density of 500 cells per well with fresh medium, growing for another 12-14 days to form colonies. Colonies are fixed with 4% paraformaldehyde (v/v), stained with 0.5% crystal violet (w/v), and counted manually. Relative formation rates were calculated, and compared using Student’s *t*-test between groups.

### Plasmids, chemicals and antibodies

All the chemicals and antibodies used in the study were summarized in Supplementary Table 1. Cell lines with stable knockdown of *EP300, DNMT1* and *DNMT3B* were obtained using lentiviral expression vector pLKO.1 (Addgene #10878) (shRNA sequences are listed in Supplementary Table 1) and were screened by puromycin (1 μg/ml). Lentivirus was packaged in HEK-293T cells. The pcDNA3.1-p300 was purchased from Addgene (#23252). The wild-type AKT1 (NM_001014431.2) was cloned in the pECMV-3×FLAG-N vector using the In-Fusion® HD Cloning Kit (Takara Bio, Cat# 639650). The K20R and K20Q mutant AKT1 plasmids were constructed using the Fast Site-Directed Mutagenesis Kit (TIANGEN, Cat# KM101). All plasmids used in this article were confirmed by Sanger sequencing.

### Pharmacological treatment of PIs

The cultured cells were treated with 1 µM MG132 for defined lengths of time (0,12, 24 and 36 h) or treated with MG132 for 24h at different concentrates (0,1,2 and 4µM). To test the effects of other PIs, cultured cells were treated with 100nM bortezomib or 10μM PI-IX for 24 hours. The control cells treated with DMSO were assessed concurrently with the experimental groups. Then cells were harvested for further assays and all experiments were repeated three times.

### Reverse transcription and quantitative PCR (RT-qPCR)

Total RNA was isolated and genomic DNA was removed by the RNA extraction kit (Vazyme, Cat# RC112-01) following the manufacturer’s instructions. Random-primed reverse transcription of RNA (1μg) was performed using Reverse Transcriptase Kit (Vazyme, Cat# R312-02). Real-time PCR analysis was performed using Green PCR Kit (TransGen, Cat# 2-21) on a QuantStudio 7 Flex system (Thermo fisher, USA). The gene-specific primers were listed in Supplementary Table 2. RT-qPCR fluorescence signal was converted to cycle times (Ct) normalized to control (ΔCt) and final results were expressed as 2^-ΔΔCt^. *B2M* or *β-actin* was used as an internal control gene for gene expression normalization.

### Western blot assay

Cellular protein extracts were prepared using total protein lysis buffer as previously reported (Du et al, 2021). Western blot assay was performed as previously described (Du et al, 2021). Briefly, protein samples were separated by 7-10% SDS-PAGE gels, and transferred to nitrocellulose (NC) membrane (Amershan protran, Cat# 10600002). The membranes were incubated with primary antibodies overnight at 4℃, and incubated with secondary peroxidase-conjugated antibodies (ZSGB-Bio company, Cat# ZB-2301, ZB-2305) for 1 hour after washing three times with TBST buffer. The blots were detected with enhanced Chemiluminescence (ECL, Advansta, Cat# 230329-20) by the ChampChemi 610 Plus System (SAGE, China). All Western blots were repeated three times independently.

### Polysome profiling assay

Polysome profiling assay was performed as previously described (Zhang et al, 2020a). Colorectal cancer cell line HCT116, either MG-132 treated or untreated, were harvested and lysed on ice with lysis buffer (20 mM Tris HCl pH 7.4, 5 mM MgCl_2_, 100 mM NaCl, 100 μg/mL cycloheximide, 1% Triton X-100, 40 U/mL RNasin and protease inhibitor cocktail). The cell extracts were centrifuged at 12,000g 4°C for 10 minutes, and then layered on a 10-50% sucrose gradient (composed of 25 mM Tris HCl pH 7.4, 5 mM MgCl_2_ and 100 mM NaCl) and centrifuged at 4°C in a SW41Ti Beckman rotor for 3.5 h at 35,000 rpm. Absorbance at 260 nm was recorded by using Piston Gradient Fractionator (Biocomp, USA). Polysomal fractions (fractions 13–22) were pooled, and RNA was isolated using the RNA extraction Kit as described above for further analysis.

### RNA-sequencing (RNA-seq) experiments and data analysis

Total RNA was extracted from cultured cells using Direct-zol RNA miniprep kit (Zymo, Cat# R2052), and ribosomal RNA was depleted using Ribo-off rRNA Depletion Kit (Vazyme, Cat# N406-01), then libraries was prepared using VAHTS Universal V8 RNA-seq Library Prep Kit for MGI (Vazyme, Cat# NRM605-01) following the manufacture’s protocols. The libraries were then sequenced on a MGISEQ-2000 sequencer (MGI) with MGISEQ-2000RS Sequencing Reagent (MGI) in paired-end 100 bp mode. For data analysis, the raw RNA-seq reads were first preprocessed to remove adapter and low-quality cycles using Ktrim software (v1.5.0) (Sun, 2020); PCR duplicates were removed using in-house programs (Zhao et al, 2024). The preprocessed reads were then mapped to human reference genome NCBI GRCh38 using STAR software (v2.7.9a) and expression quantifications were performed using featureCounts software (v2.0.3) against Ref-Seq gene annotations (Sayers et al, 2024). Differential expression analysis was performed using DESeq2 package (v1.26.0), and genes with expression changes larger than 1.5-fold and adjusted p-values lower than 0.05 were considered as differentially expressed.

### Statistical analysis

All experiments were biologically repeated at least twice, and the results were presented as mean ± SD. Mean values between two experiment-groups were compared using Student’s t-test if the data follows normal distributions, otherwise they were compared using a nonparametric test. Mean values among multiple groups were compared using one-way ANOVA. All statistical tests were two-sided, and deemed statistically significant if *p* value less than 0.05.

### Data availability

RNA-seq data has been deposited to Gene Expression Omnibus (GEO) under accession number GSE268199 (the reviewers could access the data using the token: azajwcyilvmblup), and the DNA methylation array data has been deposited to GEO under accession number GSE26851 (the reviewers could access the data using the token: krglgyscvradrwp). The data will be made publicly available upon acceptance of this manuscript.

## Acknowledgement

The study was funded by National Natural Science Foundation of China (Award Number: 32170590), Guangdong Province Department of Education (Award Number: 2021ZDZX2062), Shenzhen Science and Technology Innovation Commission (Award Number: JCYJ20220530113814033), National Key R&D Program of China (2022YFA0912700), Guangdong Basic and Applied Basic Research Foundation (2023B1515120073), Shenzhen Bay Scholar Fellowship (to C.D. and K.S.), and Major Program of Shenzhen Bay Scholars Program. We’d like to thank SZBL Supercomputing Center for computational support.

## Author Contributions Statement

**Wenwen Zhou**, **Yuling Sheng** and **Dingxue Hu:** Investigation, Data curation, Writing-review & editing, Methodology. **Yunyun An, Mengqi Yang, Wanqiu Wang** and **Jingyu Yan:** Investigation, Data curation, Methodology, Software. **Shuxia Zhang** and **Yunze Li:** Investigation, Methodology. **Shiva Basnet:** Writing-review & editing. **Qi Liu, Haoyuan Tan**: Methodology, Validation. **Yi Tan:** Supervision. **Jing Gao**: Methodology, Formal analysis. **Kun Sun**: Investigation, Methodology, Writing-review & editing, Supervision. **Changzheng Du:** Writing-original draft & editing, Funding acquisition, Conceptualization. **Wenwen Zhou**, **Yuling Sheng** and **Dingxue Hu** contributed equally to this work.

---

**Supplementary Figure 1.**
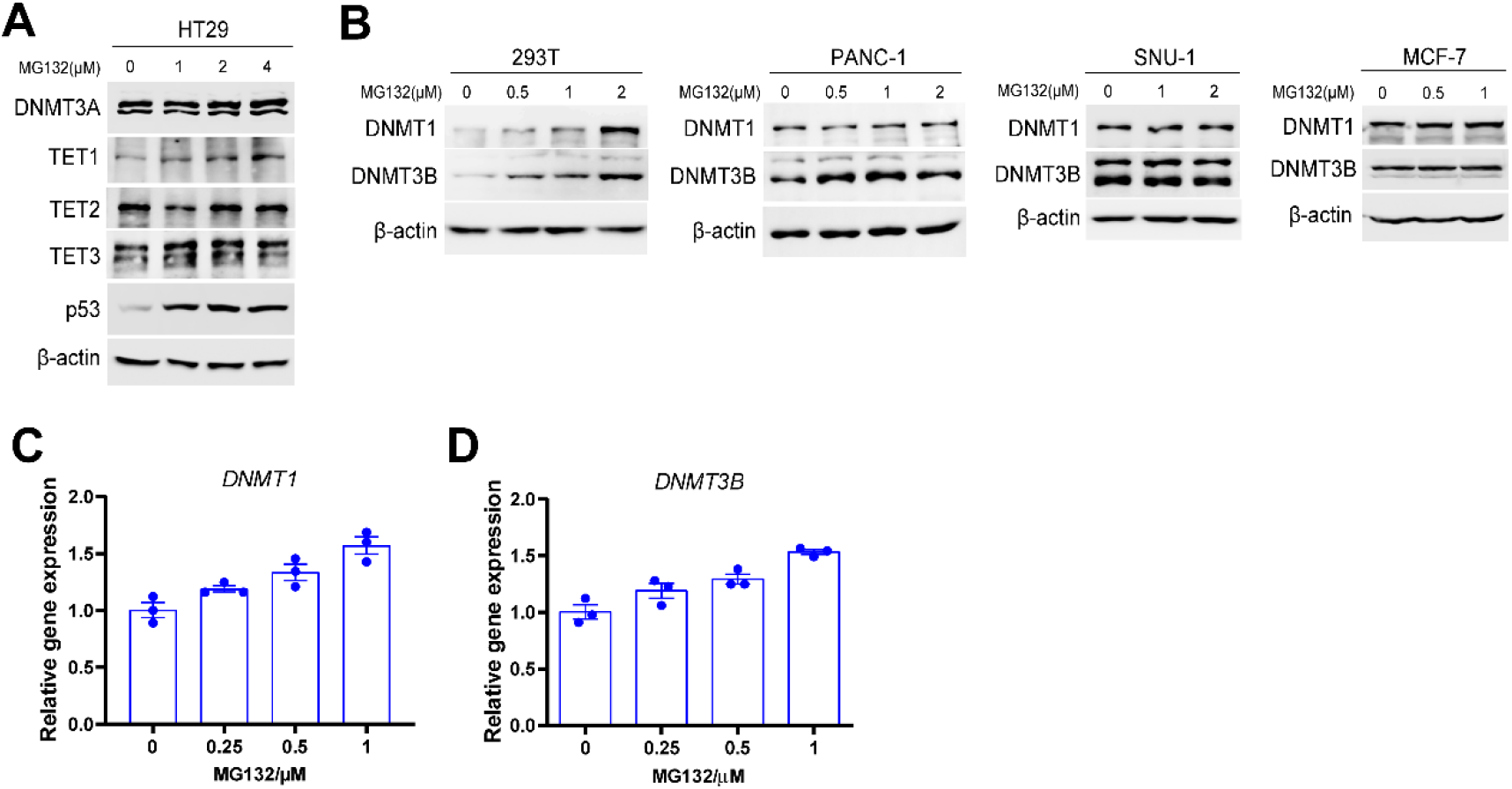
The expression of DNA methyltransferases and demethylases in multiple cell lines. **(A).** The abundance of DNMT3A, TET1, TET2 and TET3 without or with treatment of MG132 in HT29, detected by immunoblotting. **(B).** The abundance of DNMT1 and DNMT3B without or with treatment of MG132 in 293T, PANC-1, SNU-1 and MCF-7, respectively, detected by immunoblotting. **(C-D).** The mRNA abundance of DNMT1 **(C)** and DNMT3B **(D)** without or with treatment of MG132 in HT29 were determined by RT-qPCR.

**Supplementary Figure 2.**
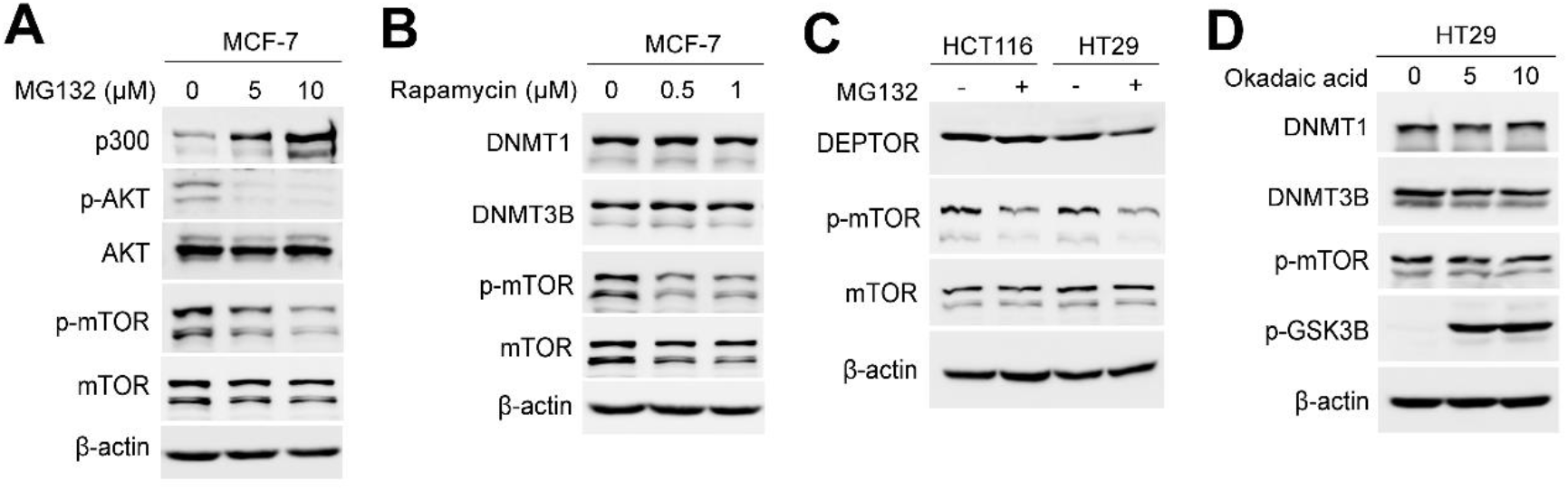
Abundance of DNMT1/3B and mTOR activity in colorectal or non-colorectal cancer cells. **(A).** The abundance of p300, p-AKT (Ser473) and p-mTOR (Ser2448) without or with treatment of MG132 in MCF, detected by immunoblotting. **(B).** The abundance of DNMT1, DNMT3B and p-mTOR without or with treatment of rapamycin, detected by immunoblotting. **(C).** The abundance of DEPTOR and p-mTOR without or with treatment of MG132 in HCT116 and HT29, detected by immunoblotting. **(D).** The abundance of DNMT1 and DNMT3B without or with treatment of okadaic acid in HT29, detected by immunoblotting.

